# Identification of novel prophages and variants of Integrative and Conjugative Elements in *Elizabethkingia anophelis* clinical isolates from Seremban, Malaysia

**DOI:** 10.1101/2024.12.10.627669

**Authors:** Asdren Zajmi, Muhamad Zarul Hanifah, Aswini Leela Loganathan, Nor Iza A. Rahman, Nurul Hafizah Mohd Yusoff, Soo Nee Tang, Qasim Ayub, Chew Chieng Yeo

## Abstract

*Elizabethkingia anophelis* is an emerging multidrug-resistant pathogen that has been identified globally, including in Malaysia. In this study, we used the Oxford Nanopore Technologies (ONT) long-read platform to generate the complete genome sequences of seven clinical isolates of *E. anophelis* from a tertiary hospital in the city of Seremban, located in the state of Negeri Sembilan in Peninsular Malaysia. These sequences were analysed alongside over 400 publicly available *E. anophelis* genomes from other countries. Three of the seven *E. anophelis* isolates, Eli4, Eli5, and Eli6, were almost identical and their isolation over a period of six weeks from the same hospital was suggestive of nosocomial transmission. Besides these three isolates, the other *E. anophelis* isolates were genetically diverse and related to distinct isolates from other countries. No plasmids were identified in the genomes of all seven *E. anophelis* isolates. A novel 39,686 bp prophage was identified in the genome of *E. anophelis* Eli4 while a presumptive incomplete prophage was found in the genomes of Eli2, Eli4, Eli5 and Eli6. *E. anophelis* genomes are notable for possessing integrative and conjugative elements (ICEs), which enable the bacterium to acquire new genes, including those encoding antimicrobial resistance or virulence. Several novel ICE sequences were discovered in the Seremban *E. anophelis* genomes, and this includes an ICE found in Eli8, designated Eli8_ICE1, which encodes phage defence genes such as type I restriction-modification systems, a bacterial cyclic oligonucleotide-based anti-phage signalling system (CBASS) and prokaryotic argonaute systems. The latter two phage defence genes have so far not been reported in the *E. anophelis* genome. This study demonstrates the utility of long-read sequencing in assessing the diversity and plasticity of *E. anophelis* genomes and emphasises further research into the role of ICEs in phage defence systems and their potential impact on antimicrobial treatment and phage therapy.

**Repositories:** The *Elizabethkingia anophelis* genomes in this study have been deposited in the NCBI database under BioProject PRJNA1175025.

## Introduction

*Elizabethkingia anophelis* is a Gram-negative bacterium which was first recognised in 2011 based on the type strain R26^T^ that was isolated from the midgut of *Anopheles gambiae* mosquitoes (Kämpfer et al., 2011; Lindh et al., 2008). The bacterium is found in both the natural environment and healthcare settings where it has been implicated in a range of infections, from mild to severe, leading to significant morbidity and mortality.

*E. anophelis* infections are more commonly reported in immunocompromised individuals and those with underlying medical conditions, such as diabetes, cancer, or chronic renal failure. *Elizabethkingia* species are naturally resistant to carbapenems, cephalosporins, aminoglycosides and most β-lactams, even in combination with β-lactamase inhibitors, due to the presence of unique intrinsic metallo-β-lactamases encoded by the *bla*_BlaB_ and *bla*_GOB_ genes, along with an extended-spectrum-β-lactamase (ESBL) encoded by *bla*_CME_ (Burnard et al., 2020).

Increasing nosocomial cases have been attributed to *E. anophelis* such as an outbreak in an intensive-care unit in Singapore in 2012 (Teo et al., 2013) and a large 2016 outbreak in Wisconsin whereby a substantial proportion of non-hospitalised patients were admitted directly from their homes (Figueroa Castro et al., 2017; Navon, 2016; Perrin et al., 2017). *E. anophelis* is commonly underreported because the symptoms in infected individuals resemble those of other common pathogens and the identification of this pathogen by traditional phenotypic methods is challenging (Lin et al., 2019). Therefore, countries with very little of no application of genome sequencing as a means of pathogen surveillance would likely have underreported incidences of *E. anophelis* infections (Zajmi et al., 2022).

Besides taxonomic conformation, whole genome sequencing provides the opportunity to delve into the genetic traits that confer resistance and virulence to *E. anophelis*. One of the hallmarks of the *Elizabethkingia* genome is its substantial genetic diversity mediated by the horizontal transfer of mobile genetic elements (MGEs) such as integrative and conjugative elements (ICEs) (Xu et al., 2019). ICEs in *E. anophelis* are diverse, and some are known to carry cargo genes encoding transcriptional regulatory and antibiotic resistance genes (Xu et al., 2019). Understanding the genetic components of MGEs in *E. anophelis* can potentially enhance our ability to predict the future efficacy of antimicrobial treatment.

To the best of our knowledge, there have not been any reports of *Elizabethkingia* genomes from Malaysia. The number of *E. anophelis* genomes initially were limited but since 2023 over 400 publicly available genome sequences, mainly from China and the USA, have been reported. With limited *E. anophelis* sampling and sequencing from other parts of the world, we risk underestimating the diversity of its genome, its carriage of MGEs including ICEs, and the full impact of its antimicrobial resistance and virulence factors. Therefore, in this study we generated complete genome sequences from seven clinical isolates of *E. anophelis* from Malaysia utilizing the Oxford Nanopore Technologies (ONT) long-read sequencing platform. We compared these to publicly available genomes to understand the global transmission of *E. anophelis* and identify its MGEs, particularly prophages and ICEs.

## Methods

### Ethical Approval

Ethical approval for the collection of Malaysian hospital isolates of *Elizabethkingia* spp. was obtained from the Malaysian Ministry of Health’s Medical Research Ethics Committee (MREC) under the National Medical Research Registry’s Protocol No. NMRR ID-21-02234-MQE (IIR), Medical Research Ethics Committee (MREC) of University of Malaya Medical Centre (UMMC) ID No: 202237-11059, and UniSZA Human Research Ethics Committee (UHREC) Approval No: UniSZA/UHREC/2024/633.

### Sample Collection

The bacterial isolates were collected between 2021 – 2022 from Hospital Tuanku Ja’afar (HTJ), the main tertiary hospital in the town of Seremban, located in the state of Negeri Sembilan, approximately 70 km southeast of Kuala Lumpur, the capital city of Malaysia. Isolates obtained from the respective hospitals’ laboratories were streaked onto blood agar and incubated overnight at 37°C. Pure colonies were then used for preliminary identification and for antimicrobial susceptibility testing.

### Identification and Antimicrobial Resistance Profiles

All purified isolates from blood agar plates were identified by the relevant hospitals’ laboratories using the VITEK 2 MS (bioMérieux) system. Antimicrobial resistance profiles were also obtained using the VITEK 2 MS system with the following panel of antibiotics: piperacillin-tazobactam (TZP), ceftazidime (CZD), cefepime (FEP), gentamicin (GEN), ciprofloxacin (CIP), trimethoprim-sulfamethoxazole (TMP-SMZ), imipenem (IMP), meropenem (MEM), ceftriaxone (CRO), and cefotaxime (CTX). There are no interpretation breakpoints for *Elizabethkingia* spp. in the CLSI guidelines (Lewis & James, 2022), therefore, the MIC susceptibility results interpreted here are presumptive, based on the CLSI breakpoints for the non-*Enterobacterales* (CLSI, 2024) as reported by Burnard et al. (2020) and Lee et al. (2022).

### Nucleic Acids Extraction

High molecular weight genomic DNA (gDNA) was extracted using the phenol-chloroform phase-separation method according to Sambrook and Russell (2006). Briefly, bacterial cells were cultured in nutrient broth at 37°C for 24 hours, collected by centrifugation, and resuspended in sterile PBS. Bacterial lysis was accomplished by adding a TLB buffer [100 mM NaCl; 10 mM Tris-Cl, pH 8.0; 25 mM EDTA, pH 8.0; 0.5% (w/v) SDS] containing RNase A (20 μg/mL) and Proteinase K (100 μL), followed by incubation at 50°C with mixing performed every 30 minutes by slowly rotating them end-to-end three times. Chloroform-isoamyl alcohol (24:1 ratio) were used for phase separation, followed by several washes to isolate the aqueous phase. DNA was precipitated using ethanol and washed with 70% ethanol. Genomic DNA was resuspended in 100 μL EB to promote DNA solubility and to prevent clumping. The final gDNA concentration was evaluated using Qubit 2.0 Flourometer (Invitrogen, Life Technologies), and a NanoDrop microvolume spectrophotometer (ThermoFisher Scientific).

### Library preparation, sequencing and raw read preprocessing

The extracted *E. anophelis* gDNA was subjected to library preparation using the SQK-LSK109 (Oxford Nanopore Technologies) and EXP-NBD104 kits, following the manufacturer’s instructions. The prepared libraries were multiplexed and loaded onto MinION flow cells (FLO-MIN111, Oxford Nanopore Technologies) for sequencing. Raw sequencing data generated from the MinION runs were base called using Guppy v5 at high accuracy configuration.

### Genome assembly, polishing and quality control

Genomic sequences were assembled with Flye (Kolmogorov et al., 2019), followed by polishing using Medaka. Assembly statistics were then assessed to ensure quality. Genomes were rotated to start from the *dnaA* position using Circlator fixstart function (Hunt et al., 2015). Genome completion and contamination assessment were determined, based on detection of single copy orthologs using the CheckM2 software (Chklovski et al., 2023).

### Taxonomy assignment and genome annotation

Taxonomic assignment of genomes was done by utilising GTDBtk (Chaumeil et al., 2022) against GTDB release 207 (Parks et al., 2020). Taxonomic assignment confirmed the identity of these genomes as *E. anophelis*. Genome annotation was conducted using bakta v1.9.3 (Schwengers et al., 2021). We also searched for antimicrobial resistance using abricate (https://github.com/tseemann/abricate) against different databases such as Resfinder (Zankari et al., 2012), PlasmidFinder (Carattoli et al., 2014), ARG-ANNOT (Gupta et al., 2014), MEGARes (Doster et al., 2020), VFDB (Chen et al., 2016), CARD (Jia et al., 2016), and NCBI AMRFinder (Feldgarden et al., 2019). Phage defence systems were identified using DefenseFinder v1.2.2 (Tesson et al., 2022).

Sequence alignment and annotation of ICE sequences were visualised using EasyFig (Sullivan et al., 2011) and clinker (Gilchrist & Chooi, 2021). Other graphs were generated using matplotlib (Hunter, 2007) and seaborn (Waskom, 2021).

### Core SNP phylogenetic tree construction

Genomes were analysed against other *E. anophelis* genomes which were accessible from NCBI GenBank and RefSeq databases. SNP calls were made against the sample genome using snippy (https://github.com/tseemann/snippy). Variant calls were then merged using the snippy-core program. The core SNP tree was constructed from merged SNP calls using Gubbins v3.1.6 (Croucher et al., 2015). The phylogenetic tree was visualised using iTOL v6 (Letunic & Bork, 2021).

### Detection of mobile genetic elements

Integrative conjugative elements (ICE) and phage sequences are part of the MGE found in previous *E. anophelis* genomes. ICE sequences were initially identified using ICEberg 3.0 (M. Wang et al., 2024). These sequences were then manually inspected by comparing genome annotations with other *E. anophelis* genomes. ICE markers were identified in candidate ICE sequences, such as VirB4 ATPase (TraG), relaxase and other type 4 secretion system (T4SS) genes. This was subsequently confirmed by identifying breakpoints where the ICE supposedly inserted into the genome. Details of these curated ICE sequences are available in **Supplementary Table 1**. Phage sequences were identified using PHASTER (Arndt et al., 2016), and the results obtained also went through manual inspection.

## Results and Discussion

### Background of the isolates and antimicrobial susceptibility profiles

Seven clinical isolates obtained from Hospital Tuanku Jaafar, Seremban, Negeri Sembilan, between November 2021 – February 2022 were identified as *E. anophelis* by the hospital’s laboratory and were chosen for genome sequencing and characterisation (**Table 1**). All isolates were obtained from blood cultures.

**Table 1:**
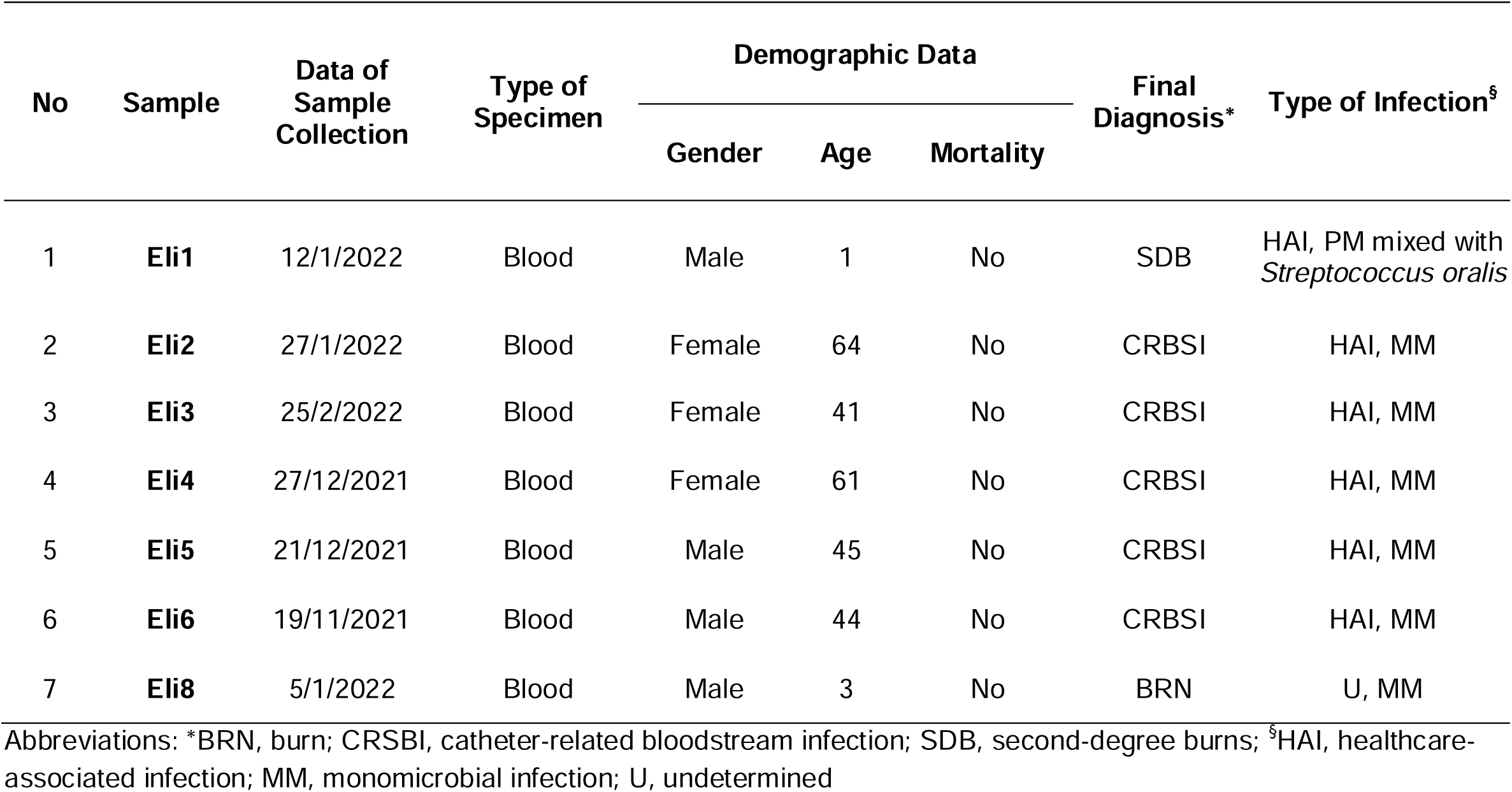
Background of the seven *Elizabethkingia anophelis* isolates from Hospital Tuanku Jaafar, Seremban, Malaysia that were sequenced in this study.

| No | Sample | Data of Sample Collection | Type of Specimen | Demographic Data |  |  | Final Diagnosis* | Type of Infection<sup>§</sup> |
| --- | --- | --- | --- | --- | --- | --- | --- | --- |
|  |  |  |  | Gender | Age | Mortality |  |  |
| 1 | Eli1 | 12/1/2022 | Blood | Male | 1 | No | SDB | HAI, PM mixed with *Streptococcus oralis* |
| 2 | Eli2 | 27/1/2022 | Blood | Female | 64 | No | CRBSI | HAI, MM |
| 3 | Eli3 | 25/2/2022 | Blood | Female | 41 | No | CRBSI | HAI, MM |
| 4 | Eli4 | 27/12/2021 | Blood | Female | 61 | No | CRBSI | HAI, MM |
| 5 | Eli5 | 21/12/2021 | Blood | Male | 45 | No | CRBSI | HAI, MM |
| 6 | Eli6 | 19/11/2021 | Blood | Male | 44 | No | CRBSI | HAI, MM |
| 7 | Eli8 | 5/1/2022 | Blood | Male | 3 | No | BRN | U, MM |

Abbreviations: \*BRN, burn; CRSBI, catheter-related bloodstream infection; SDB, second-degree burns; <sup>§</sup>HAI, healthcare-associated infection; MM, monomicrobial infection; U, undetermined

All seven *E. anophelis* isolates were resistant to imipenem and meropenem (carbapenems), and the third-generation cephalosporins ceftazidime, ceftriaxone, and cefotaxime (**Table 2**), as has been reported for most *Elizabethkingia* isolates (Burnard et al., 2020). Resistance to the fourth-generation cephalosporin, cefepime, was observed in six of the *E. anophelis* isolates whereas one isolate, Eli1, was susceptible and another, Eli3, showed intermediate susceptibility (with an MIC of 16 mg/L). All seven isolates were susceptible to piperacillin-tazobactam and trimethoprim-sulphamethoxazole whereas ciprofloxacin resistance was only observed in one isolate, Eli3. All isolates, except Eli1, showed resistance to the aminoglycoside, gentamicin (**Table 2**).

**Table 2:** Antibiotic susceptibility profiles of the seven *Elizabethkingia anophelis* isolates from Hospital Tuanku Jaafar, Seremban, Malaysia.

| Antibiotics* | Samples |  |  |  |  |  |  |  |  |  |  |  |  |  |
| --- | --- | --- | --- | --- | --- | --- | --- | --- | --- | --- | --- | --- | --- | --- |
|  | Eli1 |  | Eli2 |  | Eli3 |  | Eli4 |  | Eli5 |  | Eli6 |  | Eli8 |  |
|  | MIC | SCP<sup>§</sup> | MIC | SCP<sup>§</sup> | MIC | SCP<sup>§</sup> | MIC | SCP<sup>§</sup> | MIC | SCP<sup>§</sup> | MIC | SCP<sup>§</sup> | MIC | SCP<sup>§</sup> |
| CAZ | 256 | R | 256 | R | 256 | R | 256 | R | 256 | R | 256 | R | 256 | R |
| CIP | 1 | S | 0.38 | S | 8 | R | 0.25 | S | 0.25 | S | 0.5 | S | 0.064 | S |
| CRO | 64 | R | 64 | R | 64 | R | 64 | R | 64 | R | 64 | R | 64 | R |
| CTX | 64 | R | 64 | R | 64 | R | 64 | R | 64 | R | 64 | R | 64 | R |
| FEP | 1 | S | 64 | R | 16 | I | 32 | R | 24 | R | 24 | R | 32 | R |
| GEN | 1.5 | S | 16 | R | 16 | R | 16 | R | 12 | R | 12 | R | 16 | R |
| IPM | 16 | R | 16 | R | 16 | R | 16 | R | 32 | R | 32 | R | 16 | R |
| MEM | 16 | R | 16 | R | 16 | R | 16 | R | 32 | R | 32 | R | 16 | R |
| TMP-SMZ | 0.19 | S | 0.28 | S | 0.38 | S | 0.19 | S | 0.75 | S | 0.75 | S | 0.19 | S |
| TZP | 6 | S | 6 | S | 6 | S | 4 | S | 6 | S | 6 | S | 4 | S |

\* CAZ, Ceftazidime; CIP, Ciprofloxacin; CRO, Ceftriaxone; CTX, Cefotaxime; FEP, Cefepime; GEN, Gentamicin; IPM, Imipenem; MEM, Meropenem; TMP-SMZ, Trimethoprim-sulfamethoxazole; and TZP, Piperacillin-tazobactam; All minimum inhibitory concentrations (MICs) are in µg/mL.
<sup>§</sup> There are no recommended interpretation breakpoints for *Elizabethkingia* spp. in the CLSI (2024). The susceptibility results presented here are presumptive and based on the interpretation breakpoints for the non-*Enterobacterales*. Abbreviations: SCP, Susceptibility; S, Susceptible; R, Resistance; I, Intermediate.

### Carriage of antimicrobial resistance genes in the E. anophelis *isolates*

Using the MinION Oxford Nanopore sequencing platform, we successfully generated seven complete *E. anophelis* genomes. The genome assembly statistics are presented in **Table 3**). Genome completion and contamination were deemed suitable for downstream analyses. All seven isolates had a single circular chromosome each without any identifiable plasmids.

**Table 3:** Genome assembly statistics for the seven *E. anophelis* isolates. *E. anophelis* type strain R26 (accession no. GCA_002023665) was included for comparison.

| Sample | Eli1 | Eli2 | Eli3 | Eli4 | Eli5 | Eli6 | Eli8 | R26<sup>T</sup> |
| --- | --- | --- | --- | --- | --- | --- | --- | --- |
| Genome size (bp) | 3,963,679 | 4,039,782 | 3,859,971 | 4,238,046 | 4,265,224 | 4,265,228 | 4,065,131 | 4,058,311 |
| N50 | 3,963,679 | 4,039,782 | 3,859,971 | 4,238,046 | 4,265,224 | 4,265,228 | 4,065,131 | 4,058,311 |
| Number of contigs | 1 | 1 | 1 | 1 | 1 | 1 | 1 | 1 |
| Completeness (%) | 100 | 98.53 | 99.7 | 100 | 100 | 100 | 100 | 100 |
| Contamination | 0.63 | 1.23 | 0.46 | 0.07 | 0.09 | 0.19 | 0.11 | 0.04 |
| GC content (%) | 35.68 | 35.57 | 35.65 | 35.4 | 35.38 | 35.38 | 35.72 | 35.46 |

All seven *E. anophelis* genomes carried the three previously described β-lactamases that are intrinsic and characteristic of *Elizabethkingia* species: the metallo-β-lactamases *bla*_BlaB_ and *bla*_GOB_ that confer resistance to carbapenems and penicillin-β-lactamase inhibitor combinations, and the extended-spectrum β-lactamase *bla*_CME-1_ that confers resistance to cephalosporins and other β-lactams (Burnard et al., 2020). All isolates also harboured the tetracycline resistance gene *tet(X)*, the chloramphenicol resistance gene *catB8*, and the aminoglycoside resistance gene *aadS*, as had been reported for other *E. anophelis* isolates (Burnard et al., 2020). No other antimicrobial resistance genes were identified from the genome sequences of the isolates. The phenotypic resistance displayed by these isolates mostly matched their respective genotypes except Eli1 which was susceptible to gentamicin although the genome harboured a full-length *aadS* without any detected mutations. A similar situation was reported by Burnard et al. (2020) for the Australian *E. anophelis* isolates in resistant to gentamicin. We did not test the *E. anophelis* isolates for susceptibility to tetracyclines and, thus, were unable to assess the effect of the carriage of the *tet(X)* gene in them. However, the sequenced Australian *E. anophelis* isolates also harboured the *tet(X)* gene but were susceptible to tetracycline (Burnard et al., 2020). In a recent study from Taiwan, sequenced isolates were susceptible to tetracyclines, but they lacked the *tet(X)* gene (Lee et al., 2022). In this study, only one of the seven isolates was resistant to ciprofloxacin (Eli3 with an MIC of 8 μg/mL). In *E. anophelis*, fluoroquinolone resistance has been reported to be mediated by an amino acid substitution in GyrA (either Ser83Ile or Ser83Arg) (Jian et al., 2019; Jian Ming-Jr et al., 2018; Lee et al., 2022; Lin et al., 2018). However, a comparison of the GyrA sequence from Eli3 indicated no mutation at the Ser83 residue or other mutations within the quinolone resistance-determining region (QRNR). Interestingly, no GyrA mutations were also reported in the Australian *E. anophelis* isolates that were resistant to ciprofloxacin and levofloxacin (Burnard et al., 2020). In contrast, only one of 98 Taiwanese *E. anophelis* isolates that were resistant to ciprofloxacin/levofloxacin carried the Ser83Ile or Ser83Arg mutation in GyrA. In this particular Taiwanese isolate, the expression level of the *acrB*-encoded efflux pump showed a 12.5-fold increase, suggesting its possible role in mediating fluoroquinolone resistance in *E. anophelis* (Jian et al., 2019).

### The Seremban E. anophelis isolates are diverse

The *E. anophelis* phylogenetic tree has expanded since the one presented by Xu et al. (2019) who analysed 13 complete and 23 draft (i.e., a total of 36) genomes and categorised them into three main clusters, designated Clusters I – III. Since then, Hu et al. (2022), commendably generated 197 additional genomes collected from the Strain Library of the First Affiliated Hospital of Zhejiang University, School of Medicine and later analysed a total of 318 *E. anophelis* genomes. In this study, we analysed a total of 471 *E. anophelis* genomes for phylogenetic analysis (Fig. 1). We modelled the evolutionary history of *E. anophelis* by constructing a maximum likelihood tree based on core SNPs. Similar to previous reports, many of these genomes originated from China where a large number of sequencing studies of *E. anophelis* have been carried out (Hu et al., 2022).

**Figure 1:**
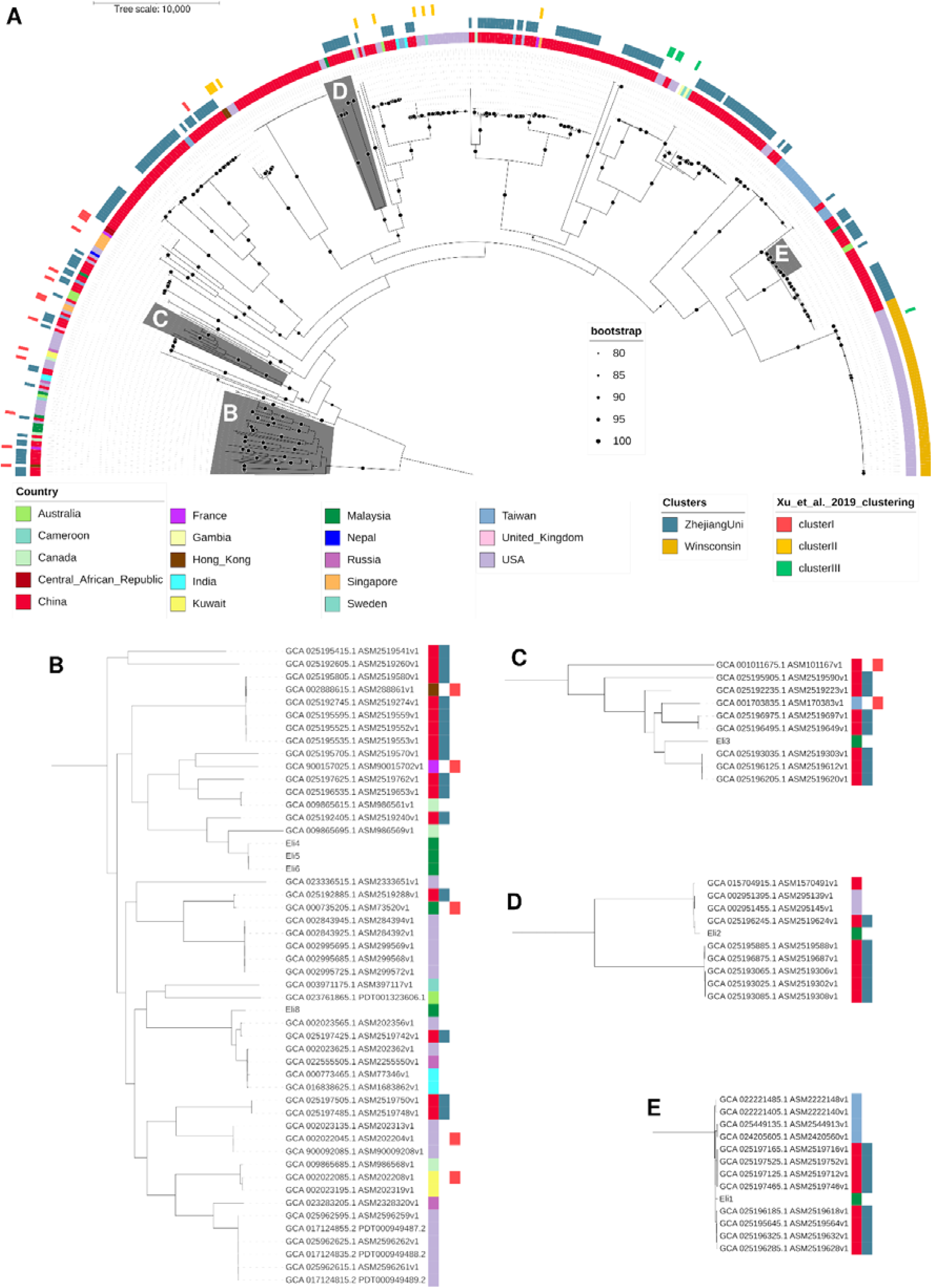
**A)** Phylogenomic tree constructed from conserved SNPs of *E. anophelis* genomes that were publicly available as of May 1, 2024. From outer to inner rings, the colours represent the three *E. anophelis* Clusters I – III categorised by Xu et al. (2019). The middle ring represents the origins of large datasets of clinical samples from the Strain Library of the First Affiliated Hospital of Zhejiang University School of Medicine, China, collected between January 2010 and April 2019 and published by Hu et al. (2022) or datasets generated from the 2015-2016 Wisconsin outbreak (Perrin et al., 2017). Innermost ring colour represents the country of origin of collected *E. anophelis*. isolates. Zoomed in selected branches of the tree are indicated in **(B) - (E)**. Note that three of the *E. anophelis* isolates, Eli4, Eli5, and Eli6, were very closely related and together with Eli8 **(B)** and Eli3 **(C)** were in Cluster I. *E. anophelis* Eli2 was in Cluster II **(D)** while Eli1 was in Cluster III **(E)**.

**Figure 2:**
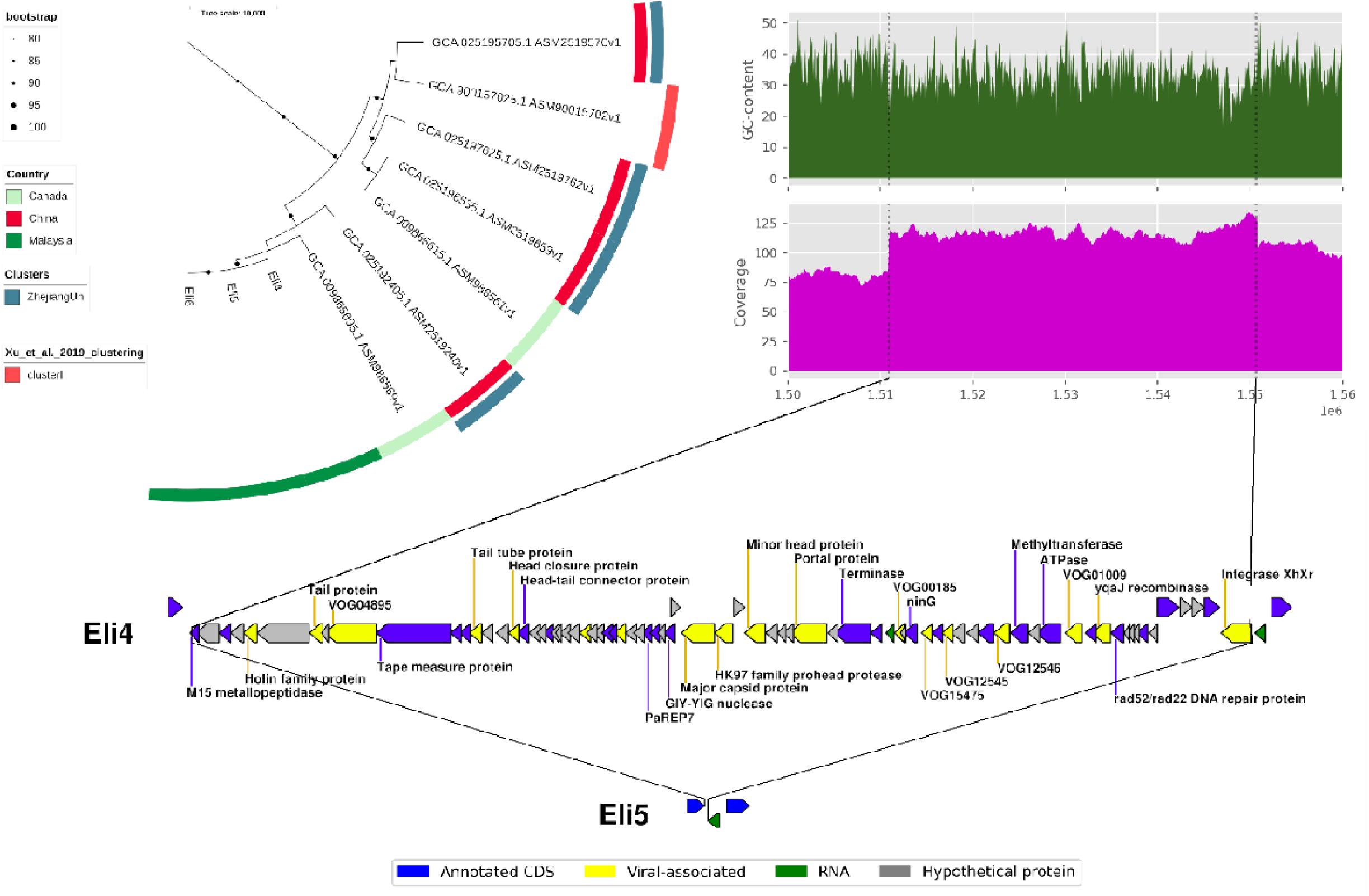
Linear genetic map of the novel prophage in *E. anophelis* Eli4, Eli4Phage1. Top left panel, subtree from Figure 1, containing *E. anophelis* Eli4, Eli5 and Eli6. Top right panel, GC content and read coverage of the Eli4 genome from nucleotides 1,510,898 to 1,550,584 which covers the Eli4Phage1 region. Bottom panel, genetic structure of Eli4Phage1 with genes and ORFs indicated by arrowheads that indicate the direction of transcription and are coloured as in the label below the figure. The prophage insertion was confirmed by absence of the sequence after aligning against its close relatives, Eli5 and Eli6.

We see isolated clusters on short branches, many of which originate from a single country, which is strongly indicative of local outbreaks. Not surprisingly, all *E. anophelis* strains from the Wisconsin outbreak in the USA in 2016 were grouped together (**Figure 1**; middle gold-coloured circle on the right), as shown previously (Hu et al., 2022). Nevertheless, some spanned across countries with most of the Cluster I isolates originating from various parts of the world. Most of the outbreak genomes were unsurprisingly generated by countries with active genome sequencing efforts. The presence of *E. anophelis* might be underestimated in other countries without a national sequencing surveillance programme.

The seven Seremban isolates were found in all three *E. anophelis* Clusters I, II and III that were designated by Xu et al. (2019) (**Figure 1**). Eli1 was found within Cluster III which included *E. anophelis* type strain R26 that was isolated from the midgut of the *Anopheles gambiae* mosquito in Stockholm, Sweden, in 2005 (Kämpfer et al., 2011) and other *E. anophelis* isolates obtained from mosquitos. Three of the Seremban isolates, Eli4, Eli5 and Eli6, were nearly identical with average nucleotide identities (ANI values) of 99.98% and above between two genomes (in comparison, these three isolates showed ANI values of between 97.4 - 98.4% with the other Seremban isolates), and were grouped together within Cluster I. These three isolates were obtained from three separate patients (one female and two males) who were diagnosed with catheter-related bloodstream infections in the months of November and December 2021 and were, thus, likely indicative of nosocomial transmission. Eli8, which was isolated in January 2022, was also within Cluster I but was grouped in a separate sub-clade (**Figure 1B**) as was Eli3 which was isolated in February 2022 (**Figure 1C**). Eli2, which was isolated in January 2022, was located within Cluster II (**Figure 1D**). Except for Eli4, Eli5, and Eli6, the remaining four Seremban *E. anophelis* isolates were diverse and distantly related to each other and were unlikely to result from nosocomial transmission.

### E. anophelis Eli4 harbours a novel prophage as well as a variant prophage that is found in other Elizabethkingia species

Despite the close relationship between Eli4, Eli5 and Eli6, we found an intact prophage sequence exclusive to Eli4, which we designated Eli4Phage1. The size of the prophage sequence is 39,686 bp, encoding 79 genes, and flanked by 15 bp direct repeat sequences (5’-TCCCTCCAGCTCCAC-3’), which were identified by PHASTER as the presumptive *attL* and *attR* sequences. Insertion of the prophage sequence was evident based on the shift in coverage within the genomic region, as well as genome alignment between Eli4 and Eli5. BLASTN results indicated that this is likely a novel prophage with no full nucleotide sequence identity coverage in the current NCBI database. Nevertheless, a 8,322 bp region of Eli4Phage1 which contained several phage proteins including the phage head-tail connector and adaptor proteins, phage tail-tube protein, and phage tape measure protein, shared 91% sequence identity with a region of the *E. anophelis* strain 0422 genome (accession no. CP016370.1), a strain which was obtained from human blood in 1950 in Florida, USA. Other regions of Eli4Phage1 only displayed nucleotide sequence identities of <1 kb with different parts of the *E. anophelis* 0422 genome.

Eli4, Eli5, and Eli6 do share a common prophage of 31,409 bp designated Eli4Phage2 which was tagged by PHASTER as an incomplete phage. Nevertheless, this presumptive prophage is flanked by 12 bp direct repeat sequences (5’-TACTTACATTTT-3’) which were identified by PHASTER as the possible *attL* and *attR* sites (**Figure 3**). Two other *E. anophelis* genomes in the database, i.e., *E. anophelis* strains 2-14 (isolated from hospital environment in Taiwan, 2015; accession no. CP071550.1), and JUNP 353 (human isolate from Nepal, 2019; accession no. AP022313.1) shared between 95 - 98% sequence identity across the entire 31,409 bp, indicating the likelihood of a similar prophage in their genomes. There are also a number of other genomes such as *E. anophelis* strains SEA01 (human blood isolate from India, 2014; accession no. CP069277.1) and NUHP1 (human isolate from Singapore, 2012; accession no. CP007547.1), *Elizabethkingia bruuniana* strain ATCC 33958 (isolated from California, USA in 1982; accession no. CP035811.1) and *Elizabethkingia endophytica* strain F3201 (human isolate from Kuwait, 1982; accession no. CP016374.1) which also shared between 95 - 98% sequence identity with Eli4Phage2 but only the first ca. 19 kb. The sequence identity spanned the presumptive *attL* site upstream of the *araJ* gene until a few nucleotides downstream of tRNA-Asn but did not cover the phage-encoded integrase, which was located 196 bp downstream of tRNA-Asn (**Figure 3**). This suggests that the Eli4Phage2 is a variant of a phage that could be found not only in *E. anophelis* but also in other *Elizabethkingia* species. Interestingly, Eli2 harbours a 28,522 bp presumptive prophage that shared nearly 100% sequence identity over a 11,283 bp region of Eli4Phage2 that spanned *araJ* to the phage_gp112-encoded gene; thereafter, the sequence identities were <95% until the phage-encoded integrase gene (a 9,801 bp region) (**Figure 3**), inferring that *E. anophelis* Eli2 likely harboured another variant of the Eli4Phage2.

**Figure 3.**
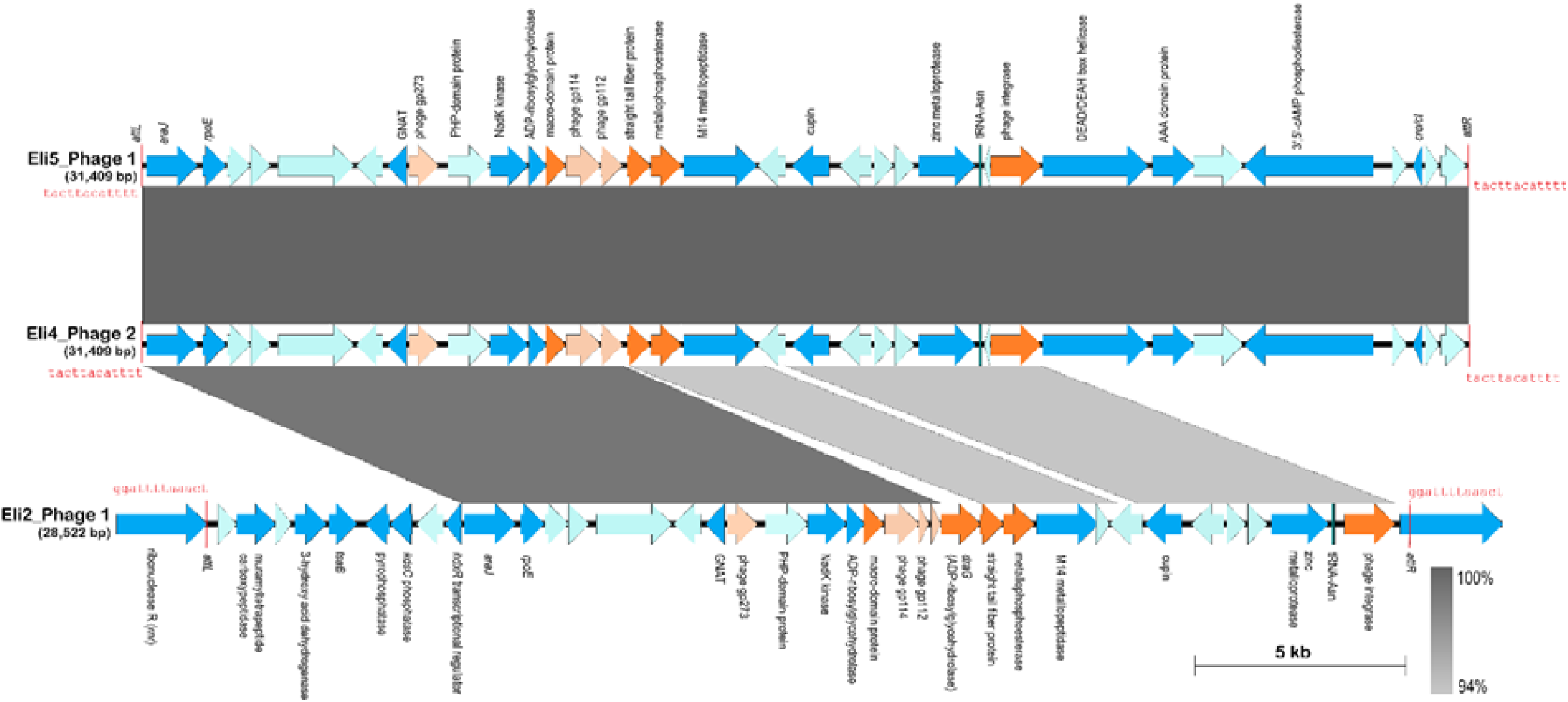
Genetic map of the Eli4Phage2 predicted by PHASTER from the genome of *Elizabethkingia anophelis* Eli4 in comparison with similar phages identified in the genomes of *E. anophelis* Eli2, Eli5 and Eli6. The Eli4_phage2 sequence (middle track) was from nts. 3,686,551 – 3,717,959 of the *E. anophelis* Eli4 genome (accession no.: SAMN44354566), the Eli5_phage1 sequence (top track) spanned nts. 3,193,129 – 3,224,537 of the *E. anophelis* Eli5 genome (accession no.: SAMN44354567), and the Eli2_phage1 sequence (Bottom track) presented here was from the reverse complement of nts. 3,435,252 – 3,406,731 of the *E. anophelis* Eli2 genome (accession no.: SAMN44354564). Note that Eli5Phage1 is identical with Eli6Phage1 and thus, only Eli5Phage1 is depicted in this Figure. The *attL* and *attR* direct repeat sequences that flank their respective prophages are indicated and shown in red-coloured fonts. Arrows show the extent and direction of the genes and ORFs. Phage-derived genes and ORFs are coloured orange with darker orange showing genes with known functions, whereas lighter orange colour indicate ORFs encoding hypothetical proteins with phage homologues. Genes or ORFs with hits in the bacterial database are shown in blue, with darker blue indicating genes with known functions while lighter blue are hypothetical proteins. The tRNA-Asn gene is indicated in green. Grey-shaded areas indicate regions with nucleotide sequence identities shown by the vertical bar at the bottom right.

The other Seremban *E. anophelis* isolates (i.e., Eli1, Eli3, and Eli8) did not harbour complete prophage sequences and only had smaller regions in their genomes (ranging from 6.4 - 16.4 kb) encoding phage-like proteins. PHASTER did not identify any potential *attL* and *attR* sites flanking these regions, indicating that these are likely partial or remnants of past prophage insertions.

### Integrative and conjugative elements (ICEs) and integrative and mobilizable elements (IMEs) in the Seremban E. anophelis genomes

One of the hallmarks of the *E. anophelis* genome is the presence of one or several integrative and conjugative elements (ICEs) which are capable of horizontal transfer between bacterial isolates via conjugation (Xu et al., 2019). ICEs typically integrate into the host chromosome and replicate along with it but sometimes they excise to form a plasmid and transfer into either another site in the genome or to another bacterial cell through conjugation (Botelho & Schulenburg, 2021; Delavat et al., 2017; Johnson & Grossman, 2015). The core components of an ICE include *attL* and *attR* (attachment sites at the left end and right end of the ICE, respectively), *int* (integrase), *xis* (excisionase), *rlx* (relaxase), *nic*/*oriT* (nicking site, origin of conjugative replication), *virD4* (ATPase/type IV coupling protein or T4CP), and either a P (TivB)- or F (TivF)-type IV secretion system (T4SS) machinery enabling mating pair formation (Delavat et al., 2017; Johnson & Grossman, 2015; Luo et al., 2023). On the other hand, integrative and mobilizable elements (IMEs) are capable of autonomous integration but are non-autonomous in conjugation since IMEs do not harbour any conjugal transfer genes and usually contain only the attachment sites *attL* and *attR*, integrase, relaxase, and the origin of conjugal transfer, *oriT* (Guédon et al., 2017).

Xu et al. (2019) had classified the ICEs identified in *E. anophelis* genomes into three types: ICEEaI (Type I), ICEEaII (Type II), and ICEEaIII (Type III). Hu et al. (2022) searched a total of 318 *E. anophelis* genomes (including 197 then-newly sequenced genomes from Zhejiang, China, along with 121 publicly available genomes) and identified 217 ICEs with the Type II ICEEaII found in majority (>100) of the isolates, followed by Type I ICEEaI (84 isolates). Thus far, no IMEs have been reported in the genomes of *Elizabethkingia.* In this study, ICEs, IMEs, or both were identified in the genomes of all seven Seremban *E. anophelis* isolates (**Table 4**). The three highly similar isolates Eli4, Eli5, and Eli6 harboured an identical ∼43.6 kb ICE, designated Eli4_ICE1, Eli5_ICE1 and Eli6_ICE1, respectively, which was inserted next to tRNA-Leu-CAA and flanked by 17-bp imperfect inverted repeats [TA(C/T)TCTTTAAATCTGTT]. Eli4_ICE1 showed sequence identities with the Type II ICE, ICEEaII(1)0422 and ICEEaII(1)CIP60.58 (which were also inserted at tRNA-Leu-CAA in their respective genomes) but these sequence identities were limited only to the *tra* genes and the genes encoding a presumptive DNA topoisomerase III (**Figure 4A**). The organisation of genes in Eli4_ICE1 suggests that it is similar to a Type II ICE, due to the presence of a RadC domain-encoding gene in between the *traD* and *traE* genes (Xu et al., 2019). However, the *virD4*/T4CP gene is contiguous with the relaxase-encoding gene as was seen in Type I ICEs and atypical of Type II ICEs, where these two genes are separated by cargo genes as seen in ICEEaII(1)0422 and ICEEaII(1)CIP60.58 (see **Figure 4A**; Xu et al., 2019). Of the genes found in Eli4_ICE1, a pair of them encode for orthologues of the Kiwa phage defence system which is typically composed of the *kwaA* and *kwaB* genes in *E. coli* (**Figure 4A**; Todeschini et al., 2023).

**Figure 4.**
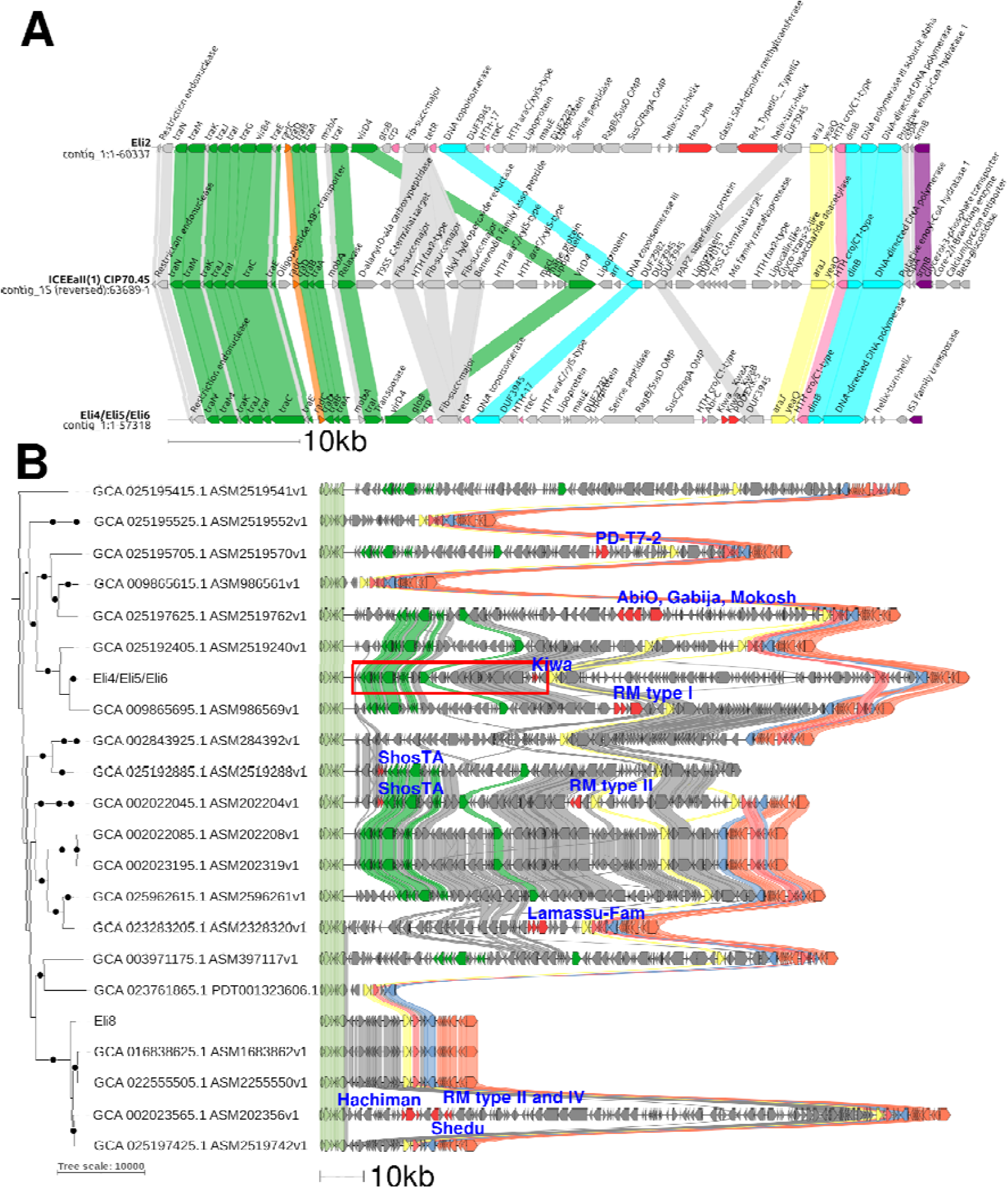
(A) Genetic map of the atypical type II ICE designated Eli4_ICE1 identified in *E. anophelis* Eli4, Eli5 (designated Eli5_ICE1), and Eli6 (Eli6_ICE1) in comparison with the reference type II ICE ICEEaII(1)_CIP60.58 (middle track). In all cases, the ICE is inserted in their respective *E. anophelis* genomes adjacent to tRNA-Leu-CAA. Arrows show the extent and direction of the genes and ORFs with T4SS genes coloured green. Orange arrows depict the gene encoding the RadC domain protein, characteristic of *E. anophelis* ICEs (Xu et al., 2019). Dark red arrows are putative phage defence genes. Light blue arrows indicate DNA polymerases and DNA topoisomerases whereas purple arrows depict helicases and transposases. Light pink arrows are putative regulatory protein-encoded genes, yellow arrows indicate the *araJ* and *yeaQ* genes. Regions with amino acid sequence identities of at least 30% are shown by linkages. (B) Comparison of genomic regions flanking Eli4_ICE1 in other *E. anophelis* genomes closely related to Eli4, Eli5 and Eli6. The maximum-likelihood phylogenetic tree of *E. anophelis* that was depicted in Figure 1 is shown on the left with Eli4 representing both Eli5 and Eli6. Eli4_ICE1 is indicated by the red rectangle with green-coloured arrows representing the T4SS genes, and the yellow-coloured arrows representing the *araJ* and *yeaQ* genes, as in **(A)**. Syntenic block of genes that were observed as conserved among most of these *E. anophelis* genomes (see **Supplementary Table 2**) are indicated as follows: light green arrows (from the 30S ribosomal protein S15-coding gene *rpsO* to tRNA-Leu), yellow arrows (for *araJ* and *yeaQ*), red arrows (for *metL1*, *qdoI*, *doxX* and a HTH-type regulator), blue arrows (for *gph*, DUF5018 and DUF2281), and orange arrows (for the lycopene cyclase-domain protein gene to *cirA*, that encodes the outer membrane receptor protein). Phage defence systems are coloured in red, with blue text annotation. Note that *E. anophelis* Eli8 does not have an ICE within this region and the ICE identified in Eli8 lies elsewhere as detailed in the text.

**Table 4:** List of ICEs and IMEs identified in the genomes of the *E. anophelis* isolates from Seremban, Malaysia.

| Genome | ICE/IME | Coordinates<br>(contig:start-end) | Size<br>(bp) | ICE Type*<br>[according to Xu et<br>al. (2019)] | Insertion<br>site | Relaxase<br>type |
| --- | --- | --- | --- | --- | --- | --- |
| Eli1 | Eli1_IME1 | contig_1:1393892-1404036 | 10,145 |  |  | MOBV |
| Eli2 | Eli2_ICE1 | contig_1:740181-789090 | 48,910 | Type II | tRNA-Leu | MOBP1 |
| Eli4 | Eli4_ICE1 | contig_1:740911-784516 | 43,606 | Type II | tRNA-Leu | MOBP1 |
|  | Eli4_IME1 | contig_1:3738276-3756769 | 18,494 |  | tRNA-Leu | MOBT |
| Eli5 | Eli5_ICE1 | contig_1:740911-784516 | 43,606 | Type II | tRNA-Leu | MOBP1 |
|  | Eli5_ICE2 | contig_1:1652533-1706153 | 53,621 | Partial Type III | tRNA-Asp | MOBP1 |
|  | Eli5_IME1 | contig_1:3263371-3263455 | 18,531 |  | tRNA-Leu | MOBT |
| Eli6 | Eli6_ICE1 | contig_1:740905-784511 | 43,607 | Type II | tRNA-Leu | MOBP1 |
|  | Eli6_ICE2 | contig_1:3204113-3257732 | 53,620 | Partial Type III | tRNA-Asp | MOBP1 |
|  | Eli6_IME1 | contig_1:1646878-1665408 | 18,531 |  | tRNA-Leu | MOBT |
| Eli8 | Eli8_ICE1 | contig_1:1774022-1819910 | 45889 | Type I |  | MOBP1 |
|  | Eli8_IME1 | contig_1:197102-226221 | 29120 |  | *bamA* | MOBC |
|  | Eli8_IME2 | contig_1:1129144-1133020 | 3877 |  | tRNA-Ser | firmi-Rep_2 |

\*As described by Xu et al. (2019); ICE type I has relaxase and *virD4* gene side by side and in opposite orientation of the rest of the other *tra* genes (refer to Figure 5). ICE type II has the RadC-domain-encoding gene between *traD* and *traE* (refer to Figure 3). ICE type III has the relaxase and *virD4* genes split by *tra* genes, all in the same orientation (refer to Figure 5). Partial ICE type III is missing a number of *tra* genes such as relaxase, *traA*, *traE*, *traG*, *traJ* and *traK* (refer Figure 3).

A majority of *E. anophelis* ICEs harbour integrases of the tyrosine recombinase (TR) type with other integrases being of the serine recombinase or the DDE-type transposase families (Xu et al., 2019) but we were unable to identify any for Eli4_ICE1. Interestingly, an almost identical ICE was found in *E. anophelis* Eli2 which was also inserted next to tRNA-Leu-CAA and flanked by identical 17 bp imperfect inverted repeats, but the designated Eli2_ICE1 was about 5 kb larger than Eli4_ICE1. Eli2_ICE1 contained additional three genes in this ∼5 kb region, two of which encode for N6-DNA methyltransferases and the other gene encode for a DEAD/DEAH box helicase (**Figure 4A**). When the ICE sequences were screened with DefenseFinder (Tesson et al., 2022), the helicase in Eli2_ICE1 was identified as an orthologue of a novel phage defence protein designated Hna that was recently characterised in *Sinorhizobium meliloti* (Sather et al., 2023). Notably *E. anophelis* Eli2 was grouped in Cluster II, which was distantly related to Eli4, Eli5 and Eli6 that were grouped in Cluster I (**Figure 1**), inferring the likelihood of past horizontal transmission of this ICE. It is possible that the ICE could have lost its integrase gene (and perhaps other genes) following its transmission and incorporation into the ancestral Malaysian *E. anophelis* genomes, or it encodes a hitherto unknown and novel integrase gene.

When comparing several genomes of *E. anophelis* that were closely related to *E. anophelis* Eli4/Eli5/Eli6 at the region surrounding Eli4_ICE1, much diversity was observed. Some genomes harbour ICEs that are similar to Eli4_ICE1 but the presence and organisation of genes within their presumptive ICEs are different when compared to Eli4_ICE (**Figure 4B**). However, in genomes such as *E. anophelis* PHOL-785 isolated in Toronto, Canada in 2018 (accession no. GCA_009865615.1) and strain SKLX065558 (GCA_025195525.1) from Zhejiang, China in 2017, the ICE was absent. In the case of the Zhejiang isolate, a large number of genes were found within this region but none of them were found to be T4SS-related (**Figure 4B**) and was, thus, not considered to be an ICE.

*E. anophelis* Eli5 and Eli6 harboured a partial Type III ICE, designated Eli5_ICE2, and which shared sequence similarities with the reference Type III ICEs ICEEaIII(7)_NUHP1, ICEEaIII(8)_NUHP1, and the partial ICEEaIII(10)_NUH6 (**Figure 5**). Type III ICEs typically harbour seven *tra* genes, i.e., *traAEGJKMN*, flanked by a relaxase and the T4CP genes (Xu et al., 2019) but in Eli5_ICE2, only *traM* and *traN* were evident along with the T4CP but without the relaxase gene (**Figure 5**). The Tyr-recombinase-type integrase was present at one end of the ICE and Eli5_ICE2 was found integrated next to tRNA-Asp-GTC, as was the case for ICEEaIII(7)_NUHP1 and ICEEaIII(8)_NUHP1, whereas the partial ICEEaIII(10)_NUH6 was integrated at tRNA-Glu-TTC (Xu et al., 2019). Both ICEEaIII(7)_NUHP1 and ICEEaIII(8)_NUHP1 were ca. 110 kb in size whereas Eli5_ICE2 was only 53.6 kb, implying that a large portion of the ICE from *traK* to the relaxase gene and accompanying cargo genes were likely lost in both Eli5 and Eli6. Interestingly, this partial Type III ICE was absent from the genome of Eli4. Among the three isolates, Eli4 was isolated on 27 December 2021 whereas Eli5 was obtained six days earlier (on 21 December 2021) and Eli6 was a full month prior (on 19 November 2021) (**Table 1**). It is thus tempting to speculate that in Eli4, acquisition of the Eli4_Phage1 (see earlier section) also led to the loss of the partial Type III ICE, Eli5_ICE2. This showcases the fluidity of the *E. anophelis* genome in these three highly related strains which were isolated within a period of about five weeks of each other within the same hospital and likely spread via nosocomial transmission.

**Figure 5.**
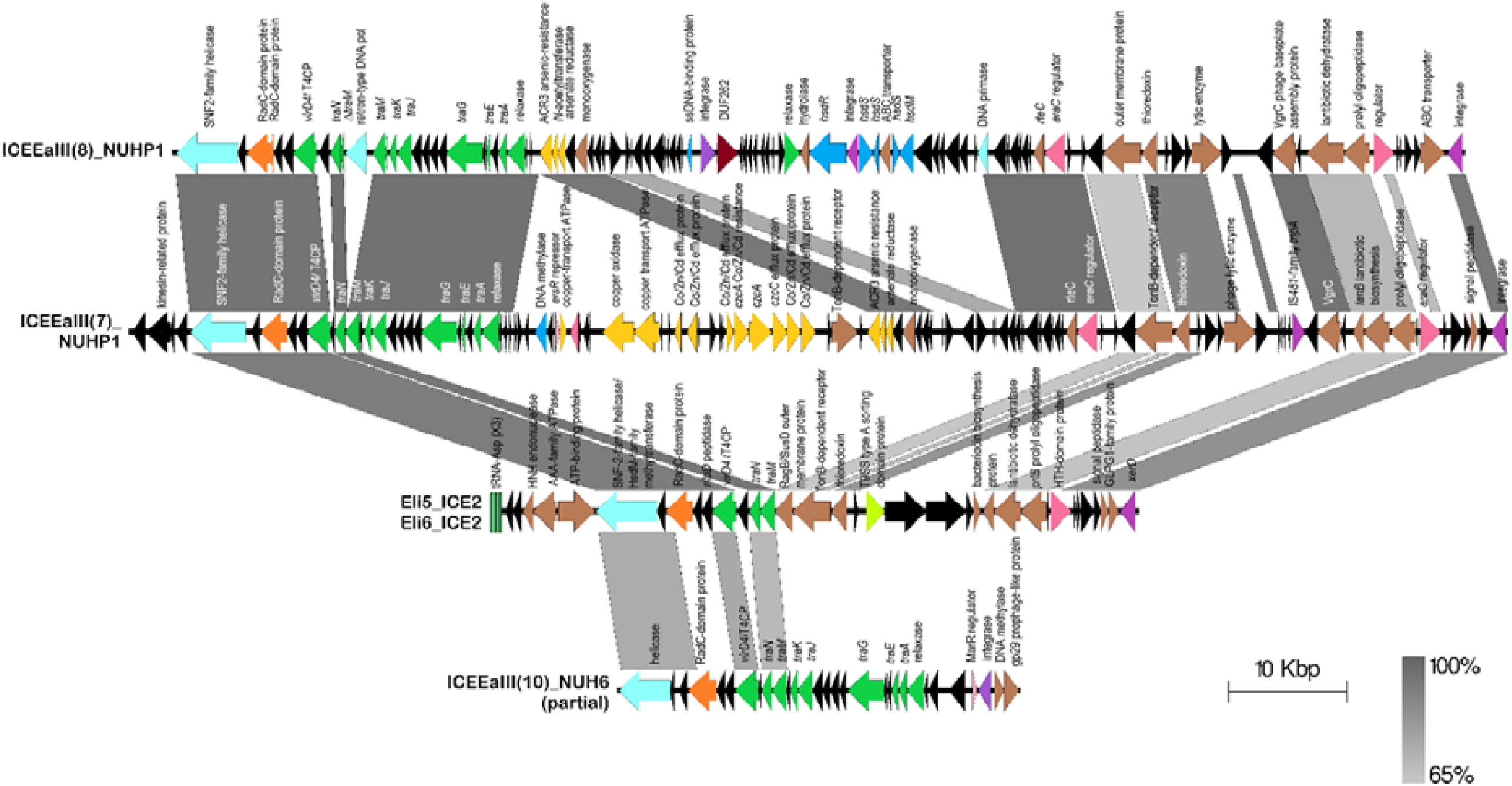
Comparative linear genetic maps of the partial Type III ICE, Eli5_ICE2 found in *E. anophelis* Eli5 (and in Eli6), with the reference Type III ICEs, ICEEaIII(8)_NUHP1 (first track) and ICEEaIII(7)_NUHP1 (second track) and the partial ICEEaIII(10)_NUH6 from Xu et al., 2019 (bottom track). T4SS genes are depicted as green arrows, orange arrows show the ORF encoding the RadC domain protein, light blue arrows indicate DNA helicases and DNA polymerases. Tyr-recombinase-type integrases are shown as purple arrows, pink arrows depict presumptive regulatory genes, brown arrows are ORFs with known function while gold-coloured arrows indicate metal resistance genes. Black arrows are hypothetical ORFs. Grey-shaded areas indicate regions with nucleotide sequence identities shown by the vertical bar at the bottom right.

All three *E. anophelis* Eli4, Eli5 and Eli6 harboured an IME of approximately 18.5 kb which encoded a Tyr-recombinase-type integrase and a relaxase. The IME, designated Eli4_IME1, was inserted near the tRNA-Leu-TAA in all three genomes and is flanked by 18-bp direct repeats (TTTTTGTACCCCAGACGG) which are the presumptive *attL* and *attR* sites. No significant sequence similarities were detected with other known *E. anophelis* ICEs and the IME encoded mainly hypothetical proteins.

### The cargo genes of the novel Eli8 ICE encode multiple bacterial defence systems

*E. anophelis* Eli8 was predicted to harbour one ICE and two IMEs (**Table 4**). The ICE in Eli8 shared sequence similarities with several type I ICE and regions in several *E. anophelis* genomes that have not been annotated as ICE. The boundaries of this 45,889 bp ICE, which we designated Eli8_ICE1, were confirmed by manual side-by-side comparisons of nearly identical *E. anophelis* genomes that are without the ICE sequence. **Figure 6** shows the full-length *E. anophelis* Eli8_ICE1 along with other similar ICE sequences aligned in juxtaposition. Analysis of the Eli8_ICE1 sequence led to several interesting observations. Firstly, even though Eli8_ICE1 shared sequence identities with several type I ICEs, the arrangement of genes differs from the typical ICEEaI whereby the *tra* cluster of genes are uninterrupted (Xu et al., 2019). In Eli8_ICE1, the *tra* genes (specifically between *traABD* and *virB4*) were interrupted by several phage defence genes including type I restriction-modification (RM) systems, and a cyclic-oligonucleotide-based anti-phage signalling system (CBASS) (**Figure 6**). RM systems have been known to occur in *Elizabethkingia* ICEs (Xu et al., 2019) but to our knowledge, CBASS has not been reported. In the CBASS antiphage defence system, a cGAS/DncV-like nucleotidyltransferase (CD-NTase) senses phage infection through the detection of viral nucleic acids and is activated to enzymatically synthesise a cyclic nucleotide immune signal that bind to effectors triggering host cell death, thereby inhibiting phage propagation (Hobbs & Kranzusch, 2024; Wang & Zhang, 2023). The CBASS system is considered the prokaryotic ancestor of the cGAS–STING antiviral pathway in animals, which similarly relies on cyclic GMP–AMP signalling (Cohen et al., 2019). This insertion of the CBASS and RM genes between the *traABD* and *virB4* genes was also observed in similar ICEs found in the genomes of two *E. anophelis* isolates from Zhejiang, China in 2022 (accession nos. GCA_025193365.1 and GCA_025195985.1), strain 12213 from Russia, 2020 (GCA_022555505.1), and strain PW2809 from Hong Kong, 2015 (GCA_001050935.1) (**Figure 6**).

**Figure 6:**
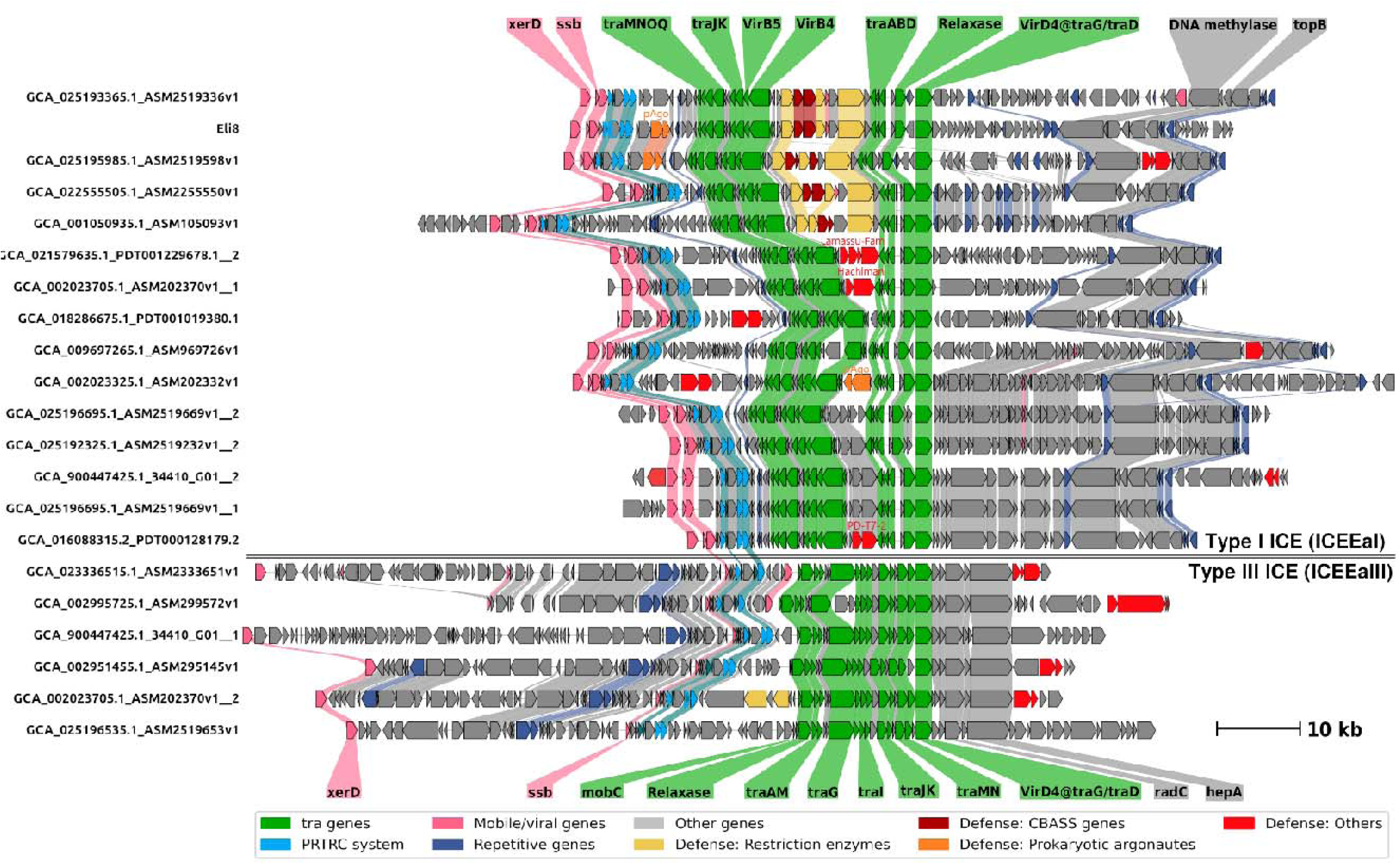
Syntenic display of different ICE structures identified across several *Elizabethkingia anophelis* genomes. Compared to the categorisation reported Xu et al. (2019), the first group corresponds to the Type I ICE*EaI*, and the second group corresponds to Type III ICE*EaIII*. Note the phage defence genes inerted between the *virB4* and *traA* genes in *E. anophelis* Eli8 (yellow arrows for restriction-modification systems; dark red arrows for the CBASS system) and related genomes. In Eli8_ICE1, and an isolate from Zhejiang, China (GCA_025195985.1) the short prokaryotic Argonaute (pAgo) genes (depicted in orange arrows) are located outside of the *tra* region whereas in an isolate from USA (GCA_002023325.1), the pAgo genes are found in between the *virB4* and *traA* genes.

Using an improved phage defence system database like DefenseFinder (Tesson et al., 2022), we found a wider diversity of phage defence systems in *E. anophelis*, especially contained in their ICE sequences. These defence systems differ greatly in structural organisation between the various *E. anophelis* ICE sequences (**Figure 6**). Besides the insertion of the CBASS and RM systems in Eli8_ICE1 and similar ICEs, other phage defence systems were also identified between the *virB4* and *traABD* genes. In the genome of *E. anophelis* SZ5977 isolated from Shenzhen, China in 2020 (GCA_021579635.1), the Lamassu-Fam phage defence genes were found in this region whereas in the genomes of two isolates obtained from the Wisconsin outbreak in the USA in 2016, this region harbours the Hachiman defence genes in GCA_002023705.1 and the PD-T7-2 two-gene system in GCA_016088315.2 (**Figure 6**). Another interesting phage defence gene that encodes a short prokaryotic Argonaute (pAgo) protein and its partner protein-coding gene was identified in between the *virB4* - *traABD* region in the ICE found in an *E. anophelis* isolate from Illinois, USA in 2016 (GCA_002023325.1). Similar pAgo and its partner genes were identified in Eli8_ICE1 and the ICE from the *E. anophelis* isolate from Zhejiang, China, 2016 (GCA_025195985.1) but they were located outside of the *tra* region and downstream of the integrase genes (**Figure 6**), inferring the likely modular nature of these genes.

Argonaute proteins are found in all domains of life, and they use small (15 - 30 bp) oligonucleotides as guides to bind and/or cleave complementary target nucleic acids and, thus, function as part of the cellular defence system (Ryazansky et al., 2018; Ugarte et al., 2023). pAgos form three large phylogenetic groups designated long-A, long-B, and short pAgos, with long-A pAgos having similar architecture to eukaryotic Argonautes (eAgos) in having an active nuclease site. However, long-B and short pAgos have substitutions in the catalytic tetrad, rendering them inactive as endonucleases. Thus, both long-B and short as nucleases, NADases, and transmembrane effectors, thereby substituting for the absence of catalytic activity in the pAgo protein (Koopal et al., 2023; Ugarte et al., 2023; C. Wang et al., 2024). Long pAgos interfere with invading plasmid and phage DNA in a guide-dependent manner, for example, the *Clostridium butyricum* Ago (CbAgo) targets both plasmid DNA and phage genomes, thereby serving as an anti-plasmid as well as an anti-phage defence module (Kuzmenko et al., 2020). In contrast, short pAgos are implicated in defence against mobile genetic elements via an abortive infection (Abi) response, utilising short RNAs as guides and triggering host cell death by mechanisms such as NAD^+^ depletion, indiscriminate cleavage of DNA and RNA, or membrane depolarisation (C. Wang et al., 2024). Interestingly, the diversity of presumptive phage defence genes found in these *E. anophelis* ICEs has not been reported before with only RM genes being indicated in some of these islands (Xu et al., 2019). However, previous research on phage defence genes in bacteria and archaea have shown that these genes tend to cluster in chromosomal regions enriched with mobile elements designated defence islands (Bernheim & Sorek, 2020; Makarova et al., 2013; Makarova et al., 2011).

Phage defence systems are frequently gained by bacteria and archaea through horizontal gene transfer but are also frequently lost from microbial genomes over short evolutionary time scales (Koonin et al., 2017), implying that there is a fitness cost in retaining these genes in the absence of selection pressure (Bernheim & Sorek, 2020). This leads to a highly variable pattern of presence and absence of these defence genes in microbial genomes (i.e., in a state of constant flux; Bernheim & Sorek, 2020) where even in closely related strains with otherwise similar genomes, the composition of defence genes can vary as indicated in the Eli8_ICE1 and its related ICEs in *E. anophelis*. It is interesting to note that no complete prophage sequence was found in the genome of *E. anophelis* Eli8, hinting at the possible functionality of these predicted phage defence genes found in Eli8_ICE1 to exclude invading phages.

Another interesting observation in Eli8_ICE1 and its related *E. anophelis* ICEs is the presence of ParB-homologs ThiF-Related Cassette (or PRTRC) systems in all of them (**Figure 6**). Very little is known about PRTRC systems and in the *E. anophelis* ICEs, they are located outside of the T4SS (i.e., *tra*) region. PRTRC systems have been found in mobile elements of other bacterial species such as plasmid pPniHBP1_2 in *Pseudomonas nitroreducens* HBP-1 (phylum Pseudomonadota) (Carraro et al., 2020) and a candidate ICE sequence in *Delftia* sp. Cs1-4 (class Betaproteobacteria) but their function is unknown (Hickey et al., 2012). Plasmid pPniHBP1_2 also carries other phage defence systems namely type-I restriction modification genes and type IV toxin-antitoxin (Carraro et al., 2020).

A putative 29,120 bp IME was detected in the Eli8 genome sequence with an protein which typically functions in lipid hydrolysis. This IME, designated Eli8_IME1, did not share any significant similarities with known *E. anophelis* ICEs, and harbours a MOBC-type relaxase, a *virB4* gene and an integrase. Eli8_IME1 also harbours an L-asparaginase, and two methyltransferase genes, one coding for a 16S rRNA methyltransferase and the other for a tRNA methyltransferase. The significance of these genes in Eli8_IME1 is currently unknown.

## Conclusion

In this study, the complete genome sequences of seven *E. anophelis* isolates obtained from a tertiary hospital in the city of Seremban, Malaysia, were presented. Three of the isolates, Eli4, Eli5, and Eli6, were closely related and likely resulted from nosocomial transmission in the hospital. The other four isolates were genetically diverse. The availability of their complete genome sequences led to the identification of novel and variant prophages and integrative and conjugative elements (ICEs). Although Eli4, Eli5 and Eli6 were closely related, Eli4 had acquired a novel 39,686 bp prophage, designated Eli4_Phage1, but had lost a partial 53.6 kb type III ICE (designated Eli5_ICE2 in Eli5 and Eli6_ICE2 in Eli6). This demonstrated the fluidity of the *E. anophelis* genomes in highly related strains that were isolated within a six-week period from the same hospital. A novel 45,889 bp variant type I ICE was discovered in the genome of *E. anophelis* Eli8 which harboured several phage defence systems. This led to the discovery of similar ICEs with other novel phage defence systems in other *E. anophelis* genomes, extending our understanding of cargo genes in ICEs that could potentially confer a protective advantage for *E. anophelis* towards bacteriophages. However, it remains to be seen if these phage defence systems could function in fending off predatory phages in *E. anophelis*, and if so, they need to be considered in any future clinical use of phage therapy for the treatment of *E. anophelis* infections. Under normal conditions, conjugation genes in ICE were not expressed (Johnson & Grossman, 2015). If phage invasion is a cue for ICE induction, this would be a good model to study the mechanisms of ICE induction, transmission and cargo gene expression.

## Author contributions

Conceptualization: A.Z., C.C.Y. Methodology: A.Z., M.Z.H., S.N.T., N.H.M.Y. Validation: Q.A., C.C.Y. Formal analysis: A.Z., M.Z.H., C.C.Y., Q.A. Resources: A.Z., Q.A., N.I.A.R., S.N.T., N.H.M.Y., C.C.Y. Supervision: C.C.Y., Q.A. Writing – original draft: A.Z., M.Z.H., C.C.Y. Writing – review and editing: Q.A., C.C.Y. Funding acquisition: A.Z., C.C.Y., Q.A.

## Ethical approvals

Ethical approval for the collection of *Elizabethkingia* hospital isolates and background clinical data was obtained from the Malaysian National Medical Research Registry and Medical Research and Ethics Committee (NMRR-MREC), Ministry of Health Malaysia, with approval number NMRR ID-21-02234-MQE (IIR), Medical Research Ethics Committee (MREC) of University of Malaya Medical Centre (UMMC) ID No: 202237-11059 and UniSZA Human Research Ethics Committee (UHREC) Approval No: UniSZA/UHREC/2024/633.

## Acknowledgments

Our thanks to the staff of the Monash University Malaysia Genomics Platform for their help in sequencing the *E. anophelis* genomes.

